# A mechanistic understanding of biofilm morphogenesis: Coexistence of mobile and sessile aggregates and phase-separated patterns

**DOI:** 10.1101/2022.06.07.494910

**Authors:** Palash Bera, Abdul Wasim, Pushpita Ghosh

## Abstract

Most bacteria in the natural environment self-organize into collective phases such as cell clusters, swarms, patterned colonies, or biofilms. The occurrence of different phases and their coexistence is governed by several intrinsic and extrinsic factors such as the growth, motion, and physicochemical interactions. Hence, it is crucial to predict the conditions under which a collective phase emerges due to the individual-level interactions. Here we develop a particle-based biophyiscal model of bacterial cells and self-secreted extracellular polymeric substances (EPS) to decipher the interplay of growth, motility-mediated dispersal, and mechanical interactions during microcolony morphogenesis. We show that depending upon the heterogeneous production and physicochemical properties of EPS, whether sticky or nonadsorbing in nature, the microcolony dynamics and architecture significantly varies. In particular, in sticky EPS, microcolony shows the coexistence of both motile and sessile aggregates rendering a transition towards biofilm formation. Wherein for the nonadsorbing EPS, which behaves as depletant in the media, a variety of phase-segregated patterned colonies either localizing the matrix component or cells at the colony periphery may emerge. We identified that the interplay of differential dispersion and the mechanical interactions among the components of the colony determines the fate of the colony morphology. Our results provide a significant understanding of the mechano-self-regulation during biofilm morphogenesis and open up possibilities of designing experiments to test the predictions.

## I. INTRODUCTION

Self-organization into multicellular communities such as swarms or biofilm-like aggregates is a common trait in most bacterial species in nature [1–11]. While developing a multicellular organization, various processes such as cell-attachment to a surface, cell growth, division, differentiation, and secretion of extracellular polymeric substances (EPS) contribute. Furthermore, cell motility and dispersion owing to physicochemical interaction among the cells and with the surrounding drives the spatiotemporal dynamics of the growing colony [12, 13]. In a microbial biofilm, many cells are embedded in a self-produced matrix of EPS containing polysaccharides or amyloid proteins or eDNA etc. [14, 15]. Biofilms formation is a route towards structural integrity, morphological diversity and protection to the cell-complex from the adverse environmental conditions. Biofilm-like aggregates cause many diseases and inflammation in animal tissues and damage in industrial applications [16, 17]. The physical interactions of the bacterial cells among themselves and with the EPS significantly influence and controls the structure and dynamics of a growing biofilms [18–22].

Over the years, several experiments have provided many insights into the individual and colony level dynamics in different bacteria [23–28]. Besides, many theoretical and computational models have developed to capture various aspects of the construction of biofilm-like structures [18, 19, 29–32]. Majority of these models followed continuum-based approaches. For example, how EPS contributes to the biofilm expansions and heterogeneous patterning are discussed in [24, 33–35] with continuum-based models. On the other hand, in a dense bacterial colony, where mechanical interactions among the bacterial cells and the surroundings appear to be a crucial factor in driving the colony expansion and morphogenesis, alternative approaches have been employed to gain insights at the mesoscopic level interactions among the components. In this context, several studies have utilized agent-based/particle-based models or a hybrid type model which can capture the individual-level interactions among the bacterial cells inside a colony [18, 19, 30, 36, 37]. Although these pre-existing studies illuminate several aspects of the occurence of collective phases, a considerable effort is needed to understand the influence of the two significant aspects of a growing bacterial colony: cell motility and physicochemical properties of self-produced EPS. In particular, how and to what extent these two aspects in conjugation with each other regulate the microcolony morphogenesis is yet to be explored.

Motility force provides individual cells a way to propel themselves in a specific direction. While, in certain conditions, a group of motile cells in a colony can exhibit long-range collective motions in the form of swirls or whirls [38–40], one of the characteristic features of developing a biofilm-like structure is the secretion and presence of extracellular polymeric substances(EPS) in a growing colony [2, 24, 34, 41–44]. From existing literature, it is now evident that the synthesis and secretion of EPS in a growing colony is a heterogeneous process [12] and it depends upon the local nutrient availability [35, 45, 46]. Apart from the heterogeneity in EPS production, the physico-chemical property of EPS can significantly control the biofilm morphogenesis. These two properties: motility and presence of EPS appear to be counteracting in a growing colony. Therefore, learning the interplay of motility and the self-produced EPS is one of the key issues which still needs considerable effort to gain insights of the spatiotemporal dynamics of a developing biofilm.

Here, we utilized a particle-based model of motile bacterial cells and self-produced EPS to study the spatiotemporal dynamics of a growing and expanding bacterial colony. We observe that depending upon the physicochemical properties of the secreted EPS, a variety of spatial morphologies can emerge in a growing multicellular colony. When the EPS is weakly attractive to the cells or sticky, we find that motile and sessile aggregates of the components coexist within the colony rendering towards the biofilm transition. The heterogeneity of EPS production across the colony profoundly impacts the coexistence of such motile and sessile cell aggregates attributing that cells do not necessarily need to be non-motile in type to develop biofilm-like structures. Instead, the heterogeneous presence of the self-produced EPS is a crucial factor. Furthermore, our results predict the spontaneous emergence of the peripheral ring of EPS when the secreted EPS is non-adsorbing to the cell surface and of higher mobility than the cells in a non-equilibrium growing colony. Interestingly, an earlier study [25] on biofilm formation reports the accumulation of EPS at the periphery but could not provide a probable mechanistic origin of the occurrence of such morphology. Our results suggest that the ratio between elasticity modulus and the friction constants is a critical controlling factor that controls the depletion effect and regulates the transition from homogeneous to a phase-segregated colony in a growing bacterial organization. The differential dispersion among the components thereby determines the outcome of colony morphology. The present study provides a systematic understanding of the mechanoregulation of microcolony morphogenesis in conjugation with cell motility and physical properties of self-secreted extracellular polymeric substances and can guide future experiments on biofilm formation.

## II. MODEL AND METHOD

We consider an individual-based model (Ghosh et.al.[19] and see *Supporting information* [SI] for details) of bacterial cells and self-secreted extracellular polymeric substances (EPS) in two dimensions. Unlike in our precedent model [19], the current study considers motile cells and EPS self-production to be dependent on the local nutrient availability as reported in the prior experimental studies [35, 45, 46]. Once the nutrient concentration *C*(*x,y*) reaches a certain level (*C*\*) it triggers EPS secretion and cells commence EPS production with a rate *k_eps_* in their surrounding medium. Moreover, the current model focuses on the physico-chemical properties of EPS where we consider two different types of EPS having different characteristics. In the first case, the self-produced EPS has a weak attraction towards the cells i.e., EPS is sticky in nature [12, 24, 45, 49, 50]. For the second type, we consider the EPS is non-adsorbing to the surface of the cells [23, 41, 47, 48] i.e., it behaves as depletants in the medium in a non-equilibrium growing colony. We will refer the aforementioned EPS as weakly-attractive or sticky and non-adsorbing or depletants throughout the text respectively. The mechanical interaction between cell-cell and EPS-EPS is repulsive (*F_rf_*) and each cell is equipped with a self-propulsion force (*F_mf_*) along the cells’ major axis. For simplicity we have put a constant term for motility force i.e *F_mf_* = *f_mot_*. When EPS is non-adsorbing in nature, we consider a repulsive mechanical force between EPS and cell particles. However, in case of sticky EPS, we considered a short range attractive force, *F_af_* between cell and EPS particles. Besides these, each particle experiences a random force *ζ*, from the surrounding medium which is taken from a uniform distribution with a range –10^-3^ to + 10^-3^ [8]. The schematic representation of the entire model is demonstrated in Figure 1.

**FIG. 1.**
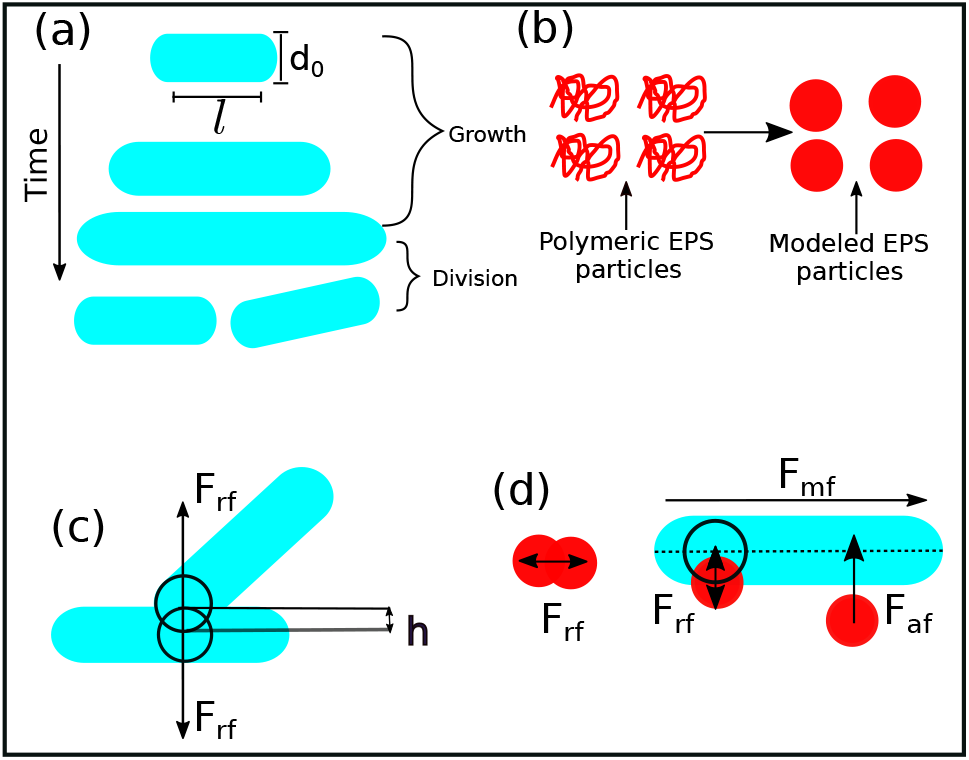
(a) A schematic representation of spherocylindrical cell with diameter *d*_0_ and cylindrical length *l.* Cells can grow as a function of time at a particular rate. It splits into two daughter cells when it reaches to a maximum length and there is a random kick in their new positions. (b) Conformation of polymeric EPS thinks about as spherical particles. (c) Repulsive mechanical interaction between two rod-shaped cells. (d) EPS-EPS and cell-EPS are interacting with a repulsive interaction if there are spatial overlaps. Cell-EPS are interacting via short-range attractive force if they are in a certain cut-off distance. Motility force acts along the long axis of the cell.

In our model, each particle follows over-damped dynamics, i.e., the medium viscosity dominates over the inertia. This suggests that the linear and the angular velocities are proportional to the force and the torque experienced by the particle respectively. Therefore, the equation of motion of each particle can be written as

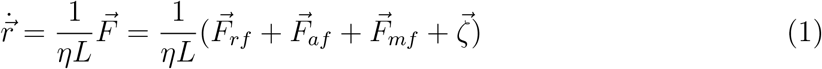

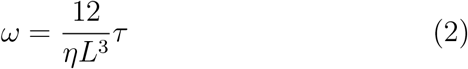

where 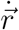, *η*, *ω* and *τ* are the linear velocity, friction co-efficient per unit length, angular velocity, and torque respectively.

The particle’s new positions and velocities are determined by solving the equations of motion using simple Euler method [51, 52] along with solving the diffusion equation for the nutrients. All parameters and constants values that are used in our simulation, are given in Table-1 in SI.

## III. RESULTS AND DISCUSSION

The present study focuses on the microcolony morphogenesis facilitated by the presence of self-produced EPS, rendering towards transitions from a colony of motile cells to biofilm-like aggregate motivated by the existing experimental studies [26]. To get a mechanistic understanding, we consider a particle-based model of bacterial cells and explicit EPS and perform computer simulations starting from a single bacterial cell developing a multicellular spatial organization. We have investigated three significant features which influence a growing bacterial colony: (i) physicochemical property of EPS: weakly attractive/sticky or non-absorbing in nature, (ii) self-propulsion force/cell motility, and (iii) nutrient-dependent cell-growth and EPS production. In response to the interplay of these factors, the morphology and spatiotemporal dynamics of a growing colony distinctly vary.

### A. Spatial organization of an expanding colony in presence of weakly attractive/sticky EPS

#### Sticky EPS facilitates biofilm transition rendering coexistence of motile and sessile aggregates

We begin our study by considering that an individual cell has self-propulsion ability; it elongates utilizing the available local nutrients and replicates when it reaches a threshold cell length. Additionally, each cell has a probability of secreting EPS in the surrounding media depending upon the local nutrient level, and those EPS are weakly attractive to the bacterial cells. A snapshot of a simulated growing colony is depicted in Figure 2(a) at a sufficiently long time (*t* = 300*h*) for cells having constant motility force *f_mot_* = 500*Pa.μm*^2^, EPS production rate *k_eps_*=1.0 hr^-1^ and for a moderate initial nutrient concentration *C*_0_ = 3.0*fg.μm*^3^. As given in Table-1 in SI, all the other parameters are kept the same throughout the text unless otherwise mentioned. The spatial organization of cells and EPS particles corresponding to Figure 2(a) is determined by calculating the radial intensity of the cells and EPS particles and depicted in Figure 2(b). The radial intensity plot shows the presence of both cells and EPS particles in the colony’s interior, whereas a thin rim of only cells retaining their motile state is observed at the periphery of the growing colony. Corresponding video (Video-S1) of the simulated colony demonstrates that cells grow, divide, move, secrete EPS in the nearby area depending upon the local accessibility of the nutrients and interact through mechanical forces to self-organize. The self-secreted EPS being a bit sticky, can self-regulate the spatiotemporal organization and movement of the components of a growing colony. The video S1 reveals the presence of apparently distinct phases of sessile aggregates and some motile cells within the colony’s interior and mobile phases at the expanding periphery. The existence of mixed phases in a growing colony attributes to biofilm morphogenesis.

**FIG. 2.**
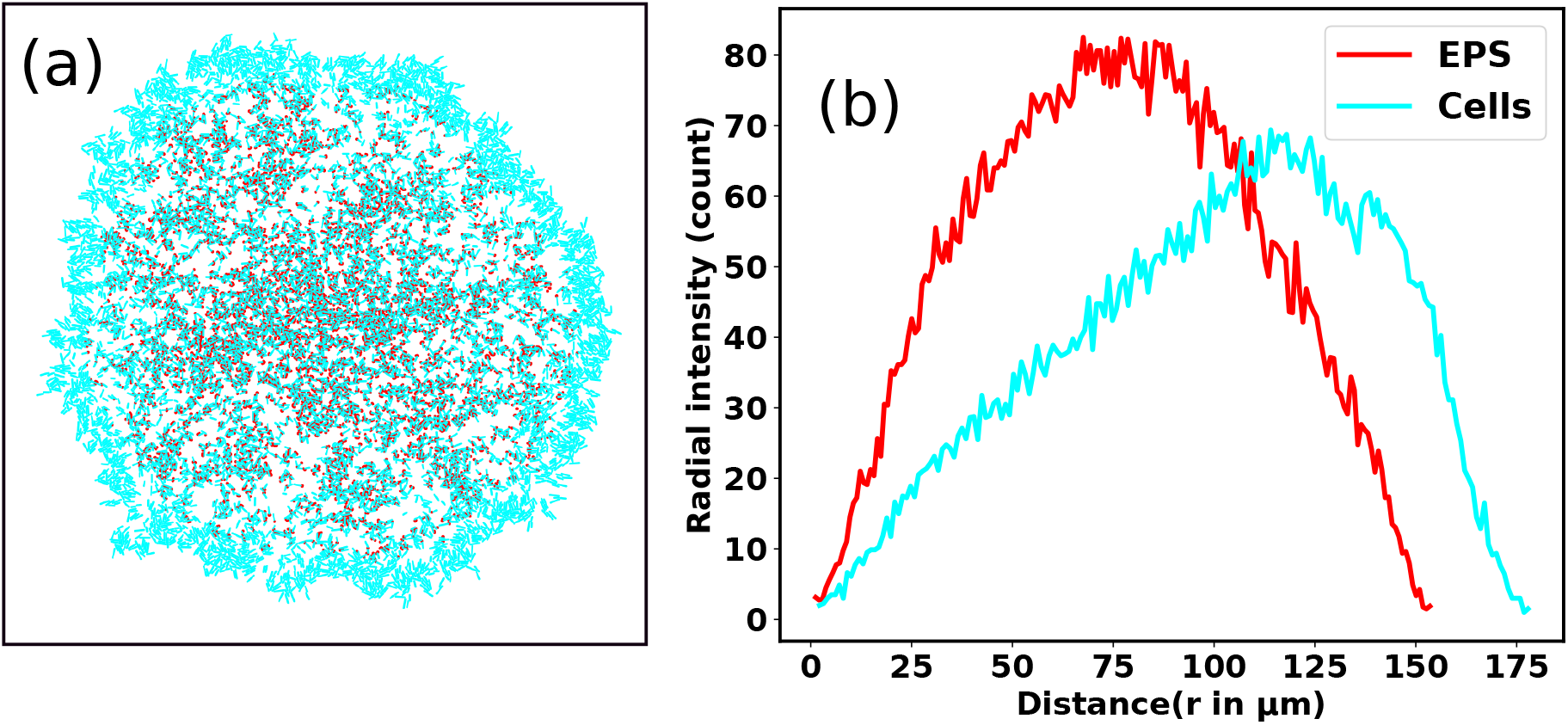
(a) A time snap of simulated colony of a growing bacteria at a long time in the presence of self-secreted sticky EPS for an initial nutrient concentration *C*_0_ = 3.0f*g.μm*^3^ and cell motility force *f_mot_* = 500*Pa.μm*^2^. Spherocylindrical bacterial cells and spherical EPS are represented in cyan and red color respectively. Peripheral cells remains in motile state, but the interior cells have initiated to transform into sessile state (almost immobilized) in response to the interaction with the sticky EPS. (b) The radial intensity profile of the cells and EPS particles as a function of the distance of the colony from the center of the simulation box determined for the Figure 2(a). EPS particles reside mostly in the interior of the colony mixed with cells and a peripheral rim of only motile cells are observed.

In order to quantify the spatial heterogeneity within the growing colony consisting both mobile and sessile aggregates, we determine the mean squared displacement (MSD) of individual cells. For dispersal of a particle, MSD is defined as 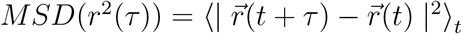, where *τ* is the lag time, 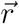 is the position of the cell, and angular brackets denote the time average. In general, MSD can be fitted with a power-law as *r*^2^(*τ*) ~ *τ^β^*, where *β* is called MSD exponent. The time profiles of MSD and MSD exponents allow us to characterize the type of dispersion i.e, how fast or slow the particles are spreading in the space. *β* = 1 indicates the standard diffusion, *β* < 1 implies the sub-diffusion, and *β* > 1 specifies the super-diffusion. Since, in a growing colony, individual bacterial life span varies with time, it is not straightforward to calculate the MSD. We strategically determine the MSD by first considering a circle about the center of the box (see *MSD calculation in SI for details*) for a particular radius. After a specific time, as the colony expands, we track those bacteria which belong to this particular radius in that period. We divide the lag time into two intervals with *τ*_1_ = (0 – 20)*h*, defined as short time lag, and *τ*_2_ = (20 – 40)*h*, defined as large time lag, to observe the different time scale behavior in observed MSD values.

Figures 3(a) and (b) demonstrate the MSD as a function of lag time for 10 cells and distribution of MSD exponents for all the cells of the colony in short lag time *τ*_1_((0 – 20)*h*) respectively. From these figures, it is clear that most of the cells show super-diffusion in a short time lag *τ*_1_. On the other hand, Figure 3(c) and (d) depict the MSD of the 10 cells for a large lag time *τ*_2_((20 – 40)*h*) which reveals the presence of both sub and super diffusion of the cells respectively. The distribution of MSD exponents of all the cells determined for large lag-time *τ*_2_ as demonstrated in Figure 3(e) also complement the fact that for a large time lag *τ*_2_; there are co-existence of cells with sub and super diffusion. As apparent from the Figure 2 and 3(e) along with the corresponding Video-S1, we observe that there are two dynamic phases of bacteria, i.e., the periphery of the colony cells are motile and at the center of the colony, they are undergoing a transition to a biofilm with the assistance of embedded EPS particles.

**FIG. 3.**
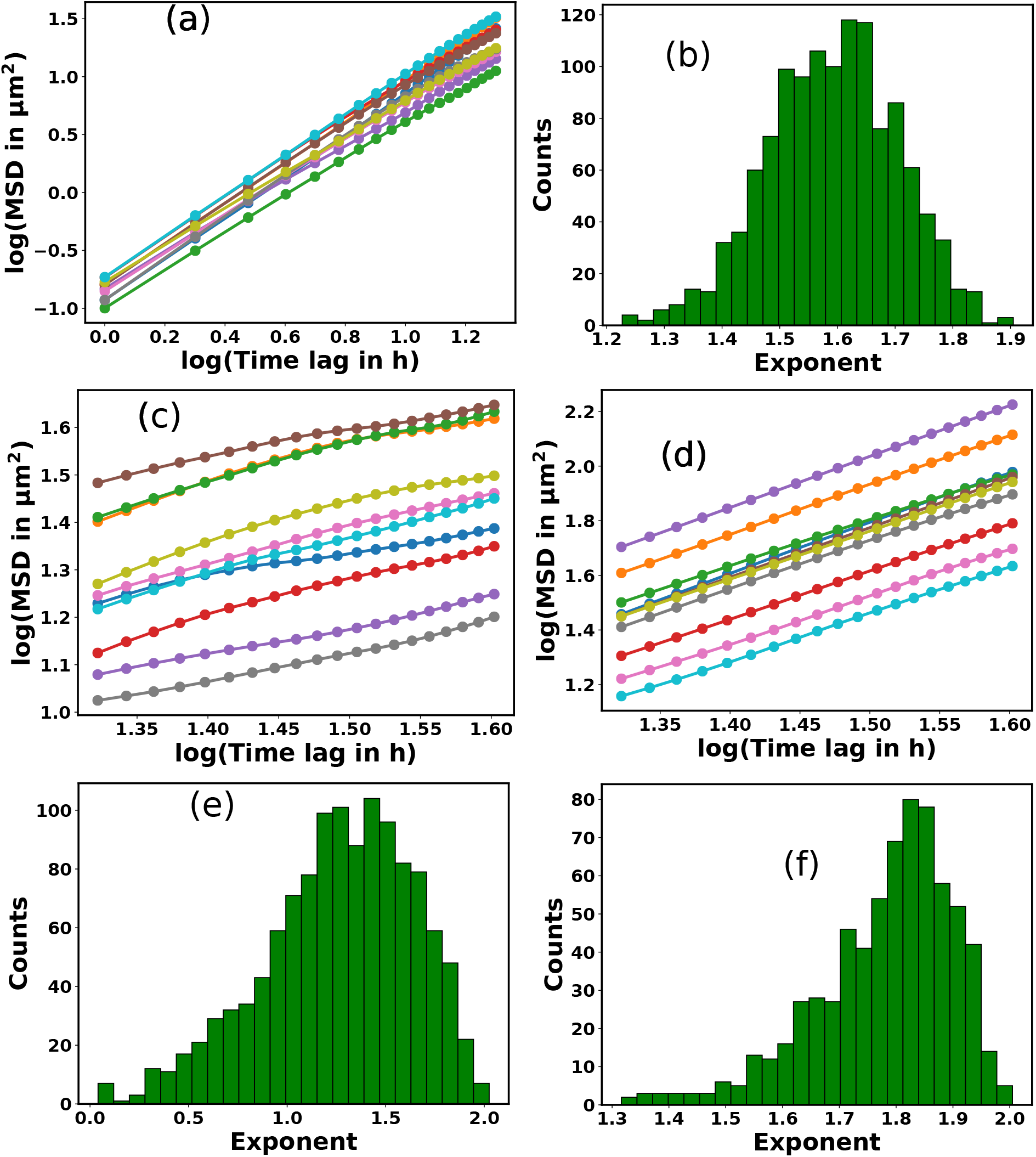
(a) MSD as a function of lag time for 10 cells from the colony and (b) the distribution of MSD exponents for all of the cells in short lag time *τ*_1_. The time profiles of MSD for (c) sub-diffusive and (d) super-diffusive cells respectively with longer lag time *τ*_2_. Here we have shown 10 trajectories for both the cases. Distribution of MSD exponents of cells in presence of (e) sticky EPS and (f) in absence EPS respectively for a longer lag time *τ*_2_. For sticky EPS, there is a co-existence of super and sub-diffusive cells. But in the absence of any EPS, almost all of the cells in a colony display super-diffusion.

At this point, we simulated a growing colony in the absence of EPS production keeping all the other parameters as same as used in Figure 2. Figure 3(f) manifests the distribution of MSD exponents for a simulated colony in the absence of EPS production for a large lag time *τ*_2_. We observe that all the bacterial cells show super-diffusion suggesting cell motility mediated fast dispersal. We extended our investigation on how EPS production controls the dispersion of the cells within the growing colony, by performing simulations for an increasing values of EPS production rate *k_eps_* as: 3.0, 5.0, 7.0, and 10.0 *h*^-1^. We evaluated the fraction of cells showing sub-diffusion by the ratio of the number of sub-diffusive cells and the total cells present within the circle (as discussed earlier). Figure 4 (a) demonstrates the bar plot of the fraction of sub-diffusive cells as a function of *k_eps_* for a particular motility force *f_mot_* = 500*Pa.μm*^2^ for a large time lag (*τ*_2_). We have also estimated the total number of EPS particles corresponding to the different values of *k_eps_* and depicted in a bar plot (Figure 4(b)). Both Figure 4(a) and (b) reveal that the fraction of sub-diffusive cells increases with *k_eps_*. However, the increase is significantly low for higher values of *k_eps_*. Since EPS production depends on several factors such as local nutrient-concentration and area density of cells as well of EPS in the two-dimensional layer, even with higher values of *k_eps_*, the EPS production will self-saturate for these restrictions. Altogether these results identify that both cell motility and heterogeneous presence of sticky EPS in a growing colony are essential for the coexistence of mobile and sessile phases rendering a biofilm transition.

**FIG. 4.**
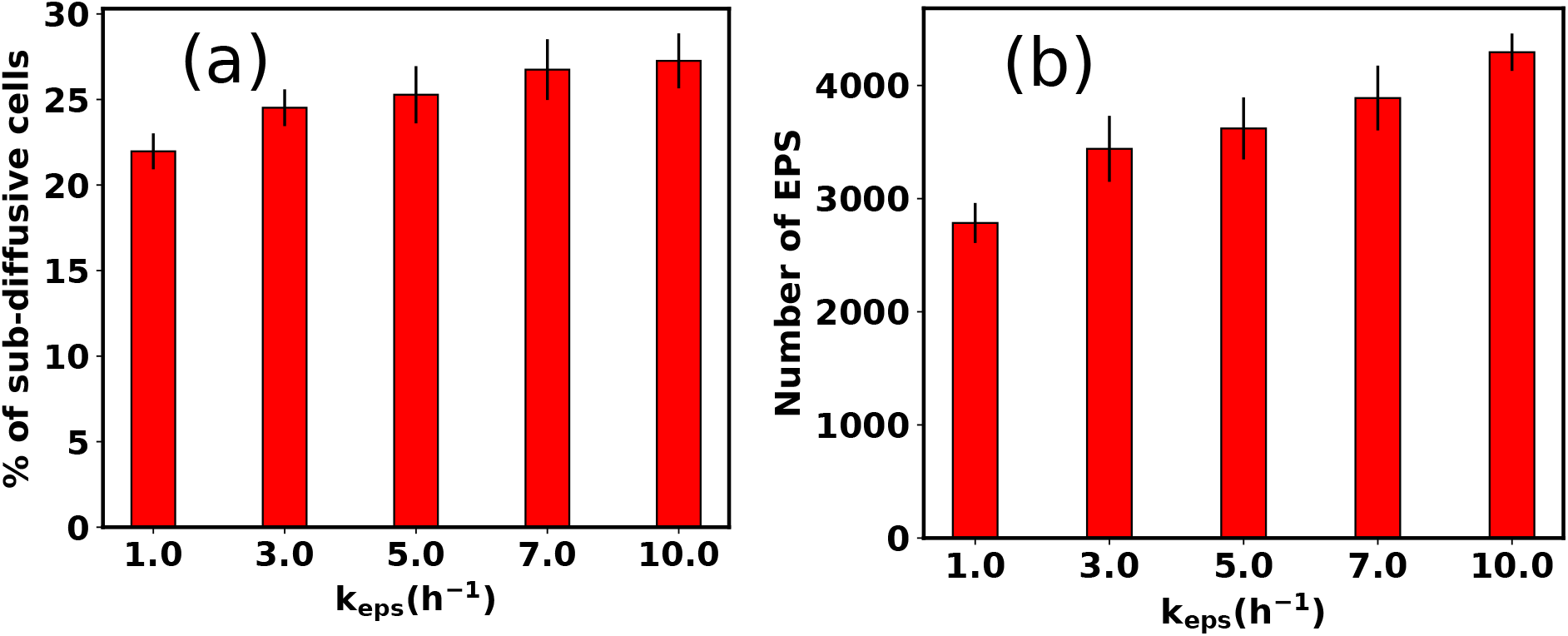
Bar plots of (a) the fraction of sub-diffusive cells and (b) the number of EPS particles as a function of EPS secretion rate (*k_eps_*).

In what follows, we will next discuss how and to what extent cell motility can regulate the colony features to get further insights into biofilm transition.

#### Cell motility regulates colony compactness and local order

For motile bacteria, self-propulsion force is a salient feature that helps them move to the nearby regions to search for nutrients for their survival. Depending on bacterial species and surrounding conditions, motility might vary. How does the variation of motility force impact the spatial architecture of a growing colony? By varying motility force, we analyze an expanding colony’s spatiotemporal organization. Figure 5(a), (b), (c), and (d) represent the snapshots of the bacterial colony at a particular time for an increasing value of self-propulsion force: *f_mot_* = 100, 300, 500, and 700 *Pa.μm*^2^ respectively, keeping all other parameters same as used in Figure 2. It is apparent from Figure 5 that the spatial morphology varies, and we observe a transition from compact to a sparse colony for the higher values of motility forces. To quantify the compactness of a growing colony, we define a quantity called “sparseness (*S_p_*)”, as 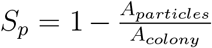, where *A_particles_* and *A_colony_* are the area of the particles and area of the colony respectively. To determine *S_p_*, we first evaluate the distance of each particle after a certain time from inoculation from the center of the simulation box and compute the maximum distance of the particle in the growing colony. We consider a circle with a radius the same as the maximum distance of the particle at that time and calculate the area of this circle and the total area of those particles belonging to this circle. So, *S_p_* ~ 1.0 indicates that particle area is very low compared to colony area, and *S_p_* ~ 0.0 suggests that particle area and colony area are almost equal. Each growing colony due to demographic noise, while simulated for the same amount of time, varies for the number of particles. We have scaled the time (min-max scaling) to compare the colony sparseness on a similar footing. Figure 5(e) illustrates the sparseness of the colony as a function of scaled time with a variation of motility forces. We observe that sparseness is high at the early stage of micro-colony morphogenesis for each colony having different motility forces. However, at the later stage, sparseness values show decay and saturate to a fixed value for each case. The sparseness value is larger for colonies with higher cell motility forces. The underlying reason of compact to sparse colony organization with increasing values of motility can be understood by considering two velocities [36]: (i) a growth-induced velocity (*v_g_*) and (ii) the self-propeling velocity (*v_mot_*) and their competition. For larger values of motility force, self-propulsion velocity *v_mot_* dominates over *v_g_*, developing more sparse colony with less order compared to the smaller values of motility force where *v_g_* dominate over *v_mot_*. Furthermore, we address how different values of motility forces can affect the colony dynamics by calculating the fraction of sub-diffusive cells as a function of *f_mot_* for large time lag (*τ*_2_) and depicted in a bar diagram as given by Figure 5(f). We observe that for *f_mot_* = 100*Pa.μm*^2^, ~ 50% of the total cells are showing sub-diffusion. But for, higher values of motility forces ~ 20% of the total cells reveal the sub-diffusive nature. This result supports that for higher values of motility forces, suppression of the mobility of cells is reduced by the self-produced sticky EPS. To shed light on how the interaction between motility and steric forces can influence the local spatial organization of the particles in a growing colony in two dimensions, we first calculate the radial distribution function of the cells. As discussed earlier, there are two types of cells within the colony; we classify them by drawing a circle from the center of mass of the colony with a slightly smaller radius compared to the maximum distance of the EPS present (see SI for details). The cells that belong to the circle are designated as the interior cells, and those outside of the circle are referred as the exterior cells. This has been done, once the colony reaches to a steady-state after a sufficiently long time. The radial distribution function is defined as 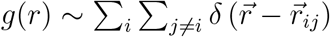, where 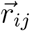 is the distance between *i^th^* and *j^th^* particle. Figure 6(a) and (b) depict the radial distribution functions of the interior and the exterior cells of the colony respectively. As demonstrated in Figure 6, there arises an initial sharp peak at a very short distance for the interior cells followed by a gradual decrease signifying an almost homogeneous colony spread of the cells. However, a close look reveals that for the interior cells, lower value of motility force (*f_mot_* = 100*Pa.μm*^2^), *g*(*r*) decays faster than the case of higher *f_mot_* = 500*Pa.μm*^2^ suggesting that motility force can affect the spatial arrangement within the interior of the colony which is undergoing a biofilm transition. For the exterior cells, the curves accompany a large peak at short distances implying formation of ordered aggregate.

**FIG. 5.**
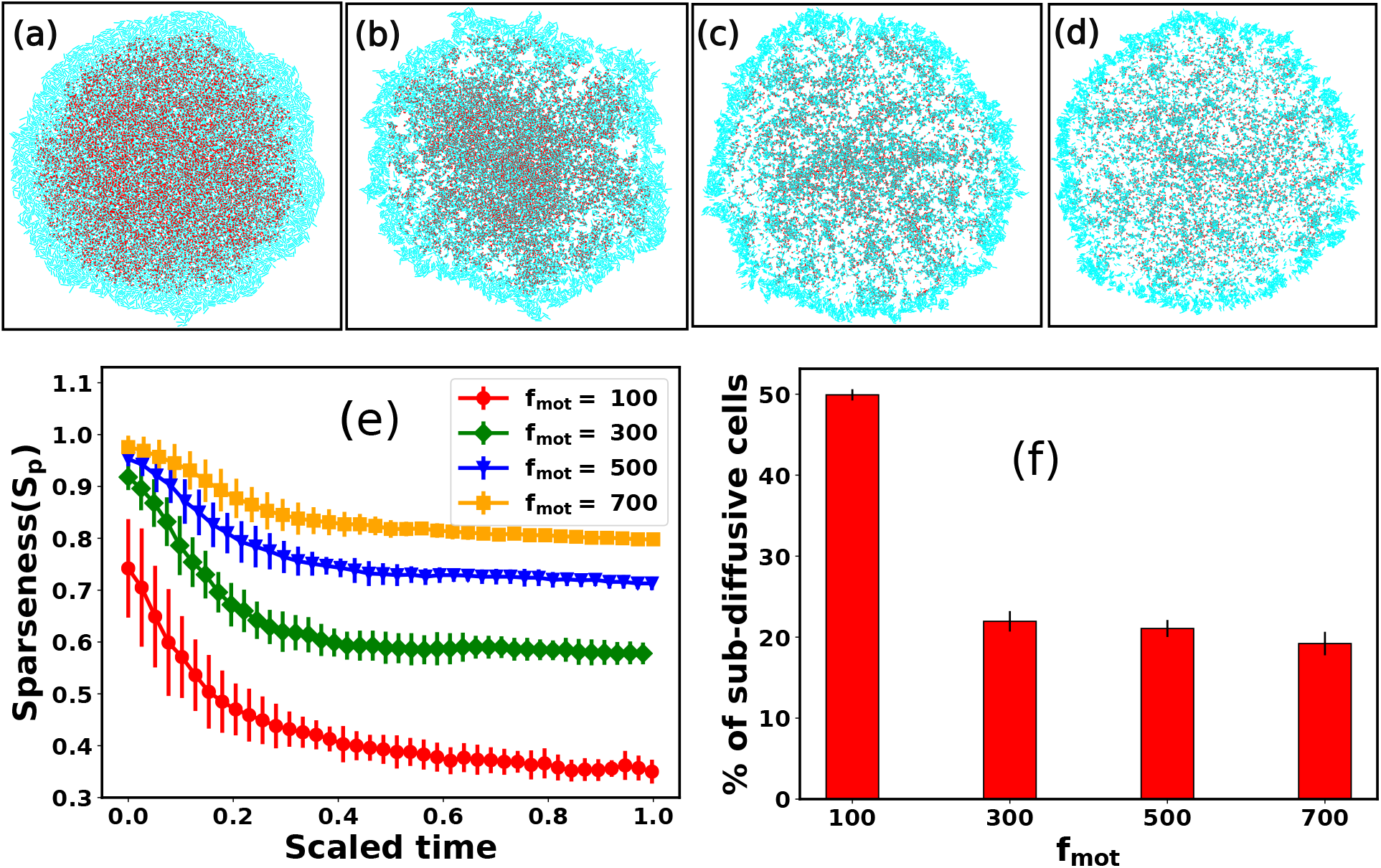
Snapshots of growing bacterial colony in the presence of sticky EPS for different values of the self-propulsion forces: (a) *f_mot_* = 100*Pa.μm*^2^, (b) *f_mot_* = 300*Pa.μm*^2^, (c) *f_mot_* = 500*Pa.μm*^2^, and (d) *f_mot_* = 700*Pa.μm*^2^ respectively. (e) Plot of Sparseness *(S_p_*) as a function of scaled time for different motility forces. (f) Percentage of sub-diffusive cells as a function of different motility forces through a bar diagram. With increasing values of motility forces, the colony becomes compact to sparse. Here error bars represent the standard error. All the other parameters remain the same as mentioned in Table-S1.

**FIG. 6.**
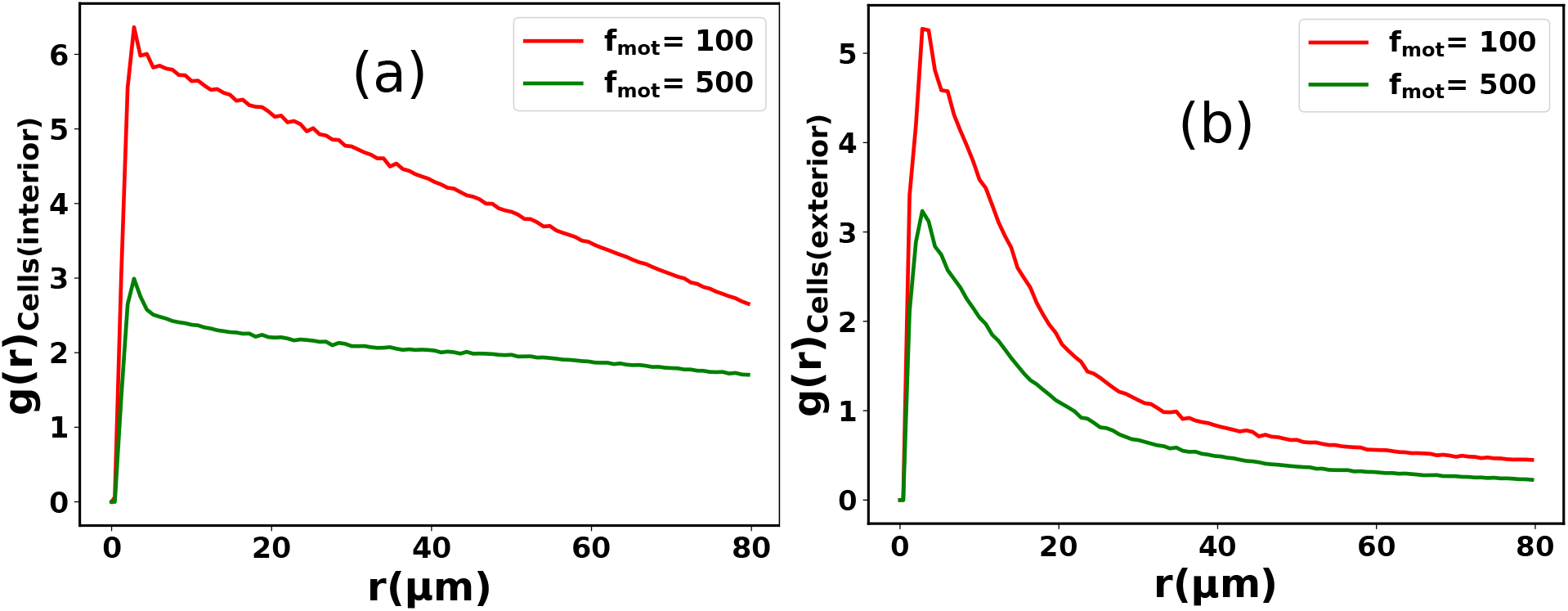
Radial distribution functions for (a) interior cells and (b) the cells at the peripheral rim of the simulated colonies, for two different motility forces: red curves correspond to *f_mot_* = 100*μm*^2^ and green curves denote the cells having *f_mot_* = 500*μm*^2^. There is a sharp peak at a short distance, for both the type of cells for different values of motility force which implies more aggregation of particles i.e compactness. All the other parameters are kept same as used in Figure 5.

To gain further insights into the local organization of the interior and the exterior cells, we compute the 2D distribution of the relative angle of cells as a function of their center of mass separation. We have calculated the relative angle of both types of cells for a particular snapshot (Figure 5) with motility force *f_mot_* = 100 and 500*Pa.μm*^2^. Figures 7(a) and (b) represent the 2d distribution of the relative angle of the interior and exterior cells respectively, as a function of their center of mass distance for motility force *f_mot_* = 100*Pa.μm*^2^. We noticed that the number of cells lying between angles of 40 to 80 is very less for exterior cells than for interior ones, which suggests that the orientation of the interior cells is relatively more random. A similar observation is observed for *f_mot_* = 500*Pa.μm*^2^ (Figure 7(c) and (d)). However, a closure look divulge that for motility force *f_mot_* = 500*Pa.μm*^2^, most of the exterior population lies between angles 0 to 60 (Figure 7(d)) whereas for motility force *f_mot_* = 100*Pa.μm*^2^ most of the exterior cells lies between angle 0 to 80 (Figure 7(b)). This observation indicates that for higher values of motility force, the exterior cells become more ordered than the lower ones.

At this stage, we will now illustrate the combined effect of the nutrient-dependent growth and motility force in colony morphodynamics.

**FIG. 7.**
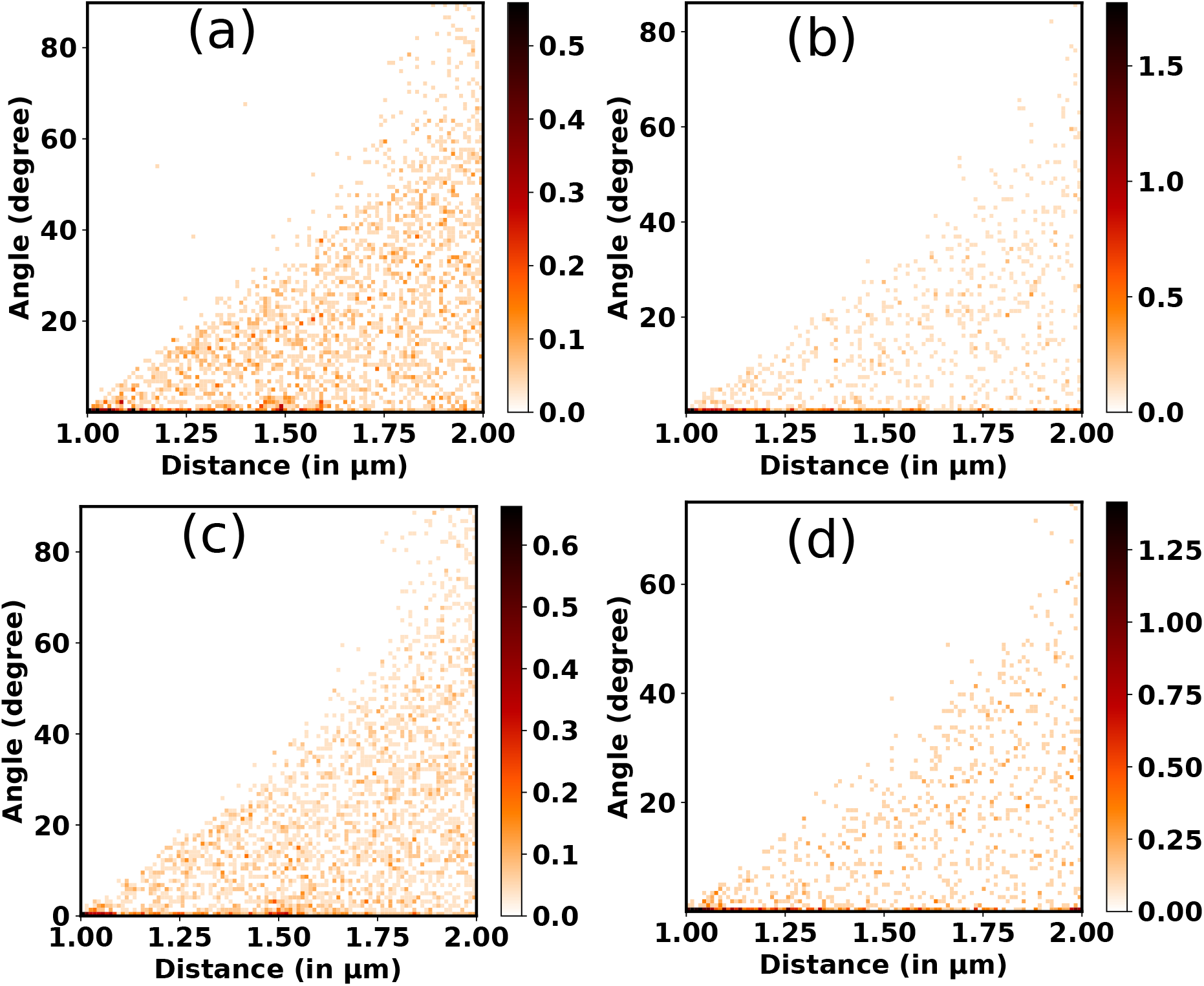
Two-dimensional (2D) distribution of angle of orientation of the (a) interior cells and (b) exterior cells with their center of mass distances for motility force *f_mot_* = 100*Pa.μm*^2^. 2d distribution of angle of orientation of the (c) interior cells and (d) exterior cells with their center of mass distances for motility force *f_mot_* = 500*Pa.μm*^2^.

#### Competition between growth-induced internal stress and motility force

In a growing bacterial colony, initial nutrient concentration *C*_0_ is a key factor that controls the growth and associated morphological dynamics. To decipher the effect of nutrient concentration, we performed computer simulations by varying *C*_0_ keeping all other parameters the same as Figure 2. Figure 8(a), (b), (c), and (d) represent the snapshots of the bacterial colony for a particular time for different values of initial nutrient concentration as *C*_0_ = 3.0, 10.0, 20.0, and 30.0*fgμm*^3^ respectively. It is apparent from Figure 8 (a)-(d) that the colony is spreading more quickly for higher values of *C*_0_. To quantify the growth dynamics concerning the variation of *C*_0_, we calculate the total area covered by the particles (cell+EPS) and the speed at which the colony expands. The front speed of the colony is determined by computing the rate of change of the colony area occupied by the particles and defining it as 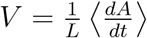, where *L* is the length of the simulation box and 〈…〉 denotes the ensemble average. Figure 8(e) and (f) illustrate the area covered by the colony and front speed as a function of time for different values of *C*_0_. For larger values of *C*_0_, cells grow faster and hit quickly to the division length and replicate, thereby causing a rapid increase in the area of the growing colony. This observation complements the higher value of the front speed of the colony.

**FIG. 8.**
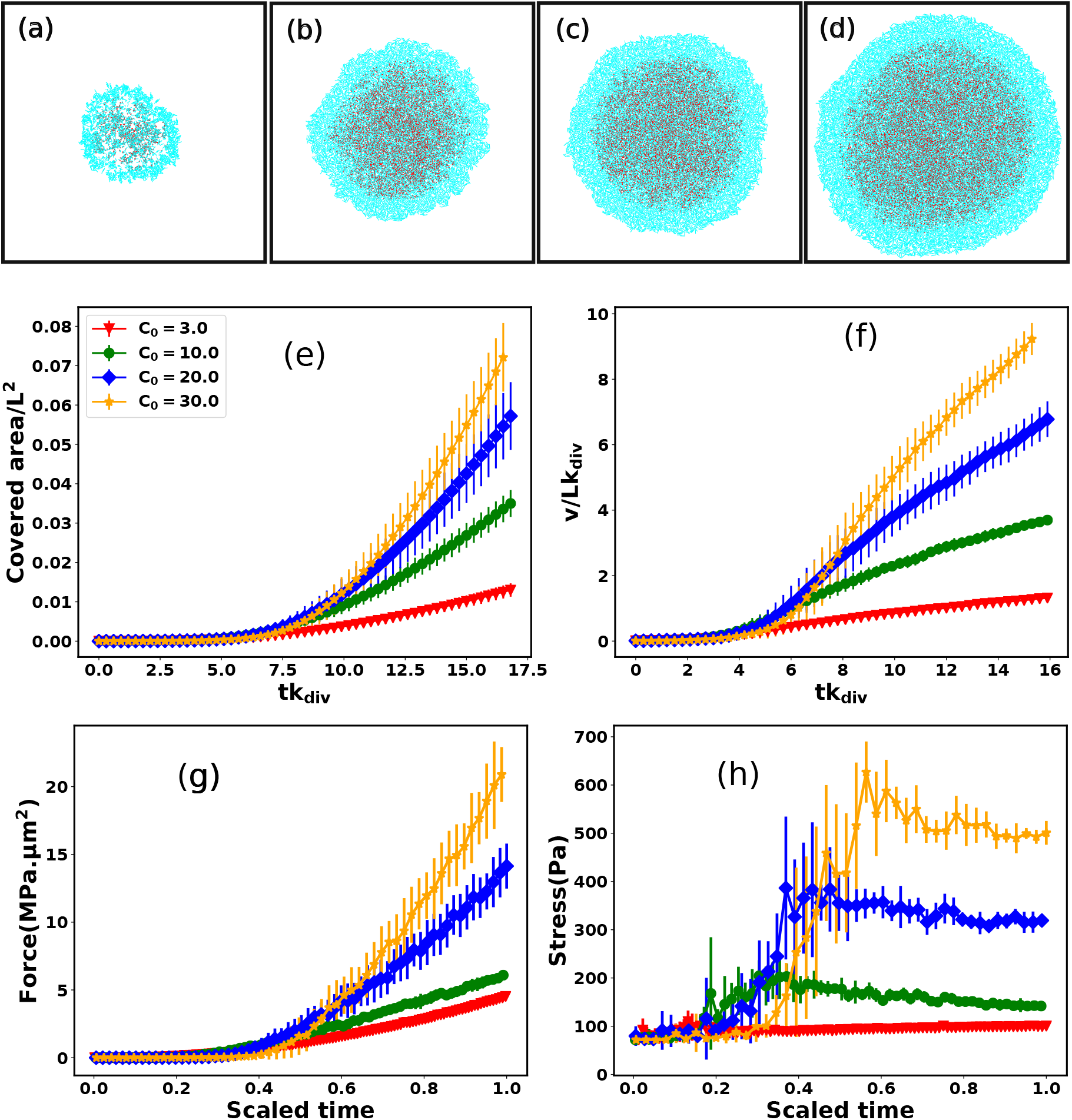
Snapshots of growing bacterial colony in presence of sticky EPS with motility force *f_mot_* = 500*Pa.μm*^2^ for different values of initial nutrient concentration: (a) *C*_0_ = 3.0*fg.μm*^3^, (b) *C*_0_ = 10.0*fg.μm*^3^, (c) *C*_0_ = 20.0*fg.μm*^3^, and (d) *C*_o_ = 30.0*fg.μm*^3^ respectively at a particular time *t* = 175*h*. With increasing values of initial nutrient concentration, the colony is spreading more quickly (e) Surface coverage and (f) front speed as a function of time for different values of initial nutrient concentration. Both are increasing with *C*_0_, due to faster cells divisions. Time profiles of (g) force and (h) stress for different values of *C*_0_. For Figures (f), (g), and (h) the color coding is the same as the legend of Figure (e). Error bar represents the standard error..

For larger values of *C*_o_, cells perceive a large mechanical force that stems from growth and rapid cell division. We evaluate total force and total stress (total force/total area), acting on the bacterial cells as demonstrated in Figure 8(g) and (h), as a function of scaled (min-max) time, respectively. The results manifest that both the force and stress are larger for a higher value of *C*_o_. However, at a longer time, the stress curve saturates. The underlying reason is that for a longer time, due to fast cell growth and divisions, the rate of change of the area of the colony is almost equal to the rate of change of total force acting on the cells leading to a saturation of the total stress. This observation suggests a competition between growth-induced internal stress and self-propulsion force which diminishes the unidirectional motion of cells. To justify this argument, we compute the MSD of cells in a growing colony for initial nutrient concentration *C*_0_ = 10.0*fg.μm*^3^ and make a distribution of MSD exponents. As depicted in Figure S1(a) and S1(b), more cells show sub-diffusion for both small (*τ*_1_) and large (*τ*_2_) lag times in comparison to *C*_0_ = 3.0*fg.μm*^3^ (Figure 3(b) and 3(e)). Due to the larger access to local nutrient concentration, cells grow, divide, and cover a certain area faster than the lower *C*_0_.

We now analyze how does the nutrient concentration profile change for different values of motility forces. We evaluate the mean value of nutrient concentration as denoted by *C_avg_* = *C*(*t*)/*C*(0), where *C*(*t*) and *C*(0) are the time-dependent and initial nutrient concentration in each grid point respectively. Figure 9 illustrates the change of *C_avg_* as a function of scaled time (*tk_div_*) for different values of motility forces. We notice that *C_avg_* of the different colonies show a similar decay with respect to the scaled time initially (up to *tk_div_* ~ 15) and differ in a later stage, showing a slower decay for the low motile cells compare to higher motile cells. For larger values of self-propulsion forces, as cells spread rapidly across the colony, they can utilize the available nutrients to grow fast and divide and multiply in numbers, thereby causing a quicker depletion of local nutrients. This behavior of colonies with different cell motility suggests that although high motility forces help in spreading in search of food, colonies with lower values of motility forces of the cells will survive for a longer time with the conserved initial nutrients.

**FIG. 9.**
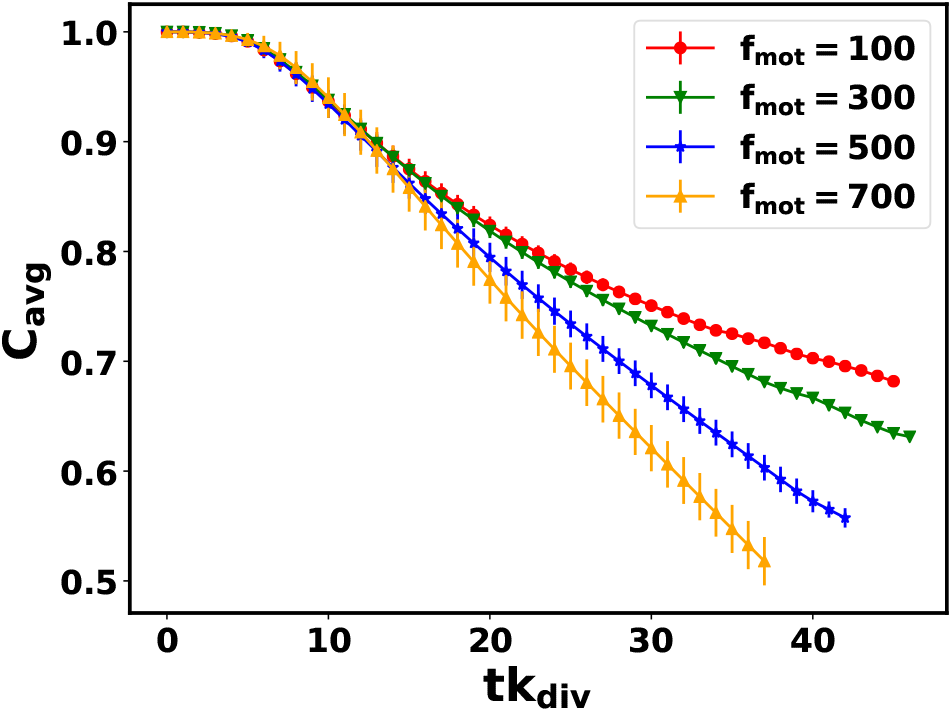
Mean nutrient concentration *C_avg_* as a function of scaled time for different values of motility forces. Here error bars represent the standard error.

So far, we have underpinned the sources of structural heterogeneity that emerge during biofilm morphogenesis. While heterogeneous growth and production of cells and EPS are crucial, it turns out that at the same time, cell motility and the weakly attractive nature of the self-secreted EPS are the driving factors towards the coexistence of mixed phases in a growing colony. The presence of both the phases is a signature of biofilm formation. Depending upon the interplay of these features as mentioned above, the colony compactness, the coexistence of motile and sessile aggregates, and their spatial organization can vary. It is essential to dig into further how and to what extent the properties of EPS might influence and control a growing colony’s spatial organization and characteristics. Existing studies report that the bacteria *S. meliloti* overproduces succinoglycan, an exopolymer, during their growth and colony expansion. Entropy-driven depletion effect contributes towards biofilm patterning as reported in [41]. In what follows, inspired by this experiment, we will now discuss the role of nonadsorbing EPS in a growing colony on a semi-solid agar surface during biofilm morphogenesis.

### B. Spatial organization of growing colony in presence of non-adsorbing EPS

#### Differential mobility facilitates localization of EPS at the peripheral region

Taking a leap from our previous study [19], here we further investigate how cell motility and non-adsorbing EPS together might impact on the spatial morphology of an expanding colony in the presence of differential frictions between the components. Since, differential friction of the components is able to cause disparity in the mobilities of the particles on hard agar surface, we have carried out simulations of growing bacterial colonies in the presence of self-secreted nonadsorbing EPS having low friction coefficient in comparison to the cells (*η_eps_* = 100*Pa.h* and *η_cell_* = 200*Pa.h*), for *C*_0_ = 3.0*fg.μm*^3^. Since, entropically driven depletion-attraction can substantially affect the morphological dynamics of a nonequilibrium growing colony in presence of non-adsorbing EPS particles [19], we vary the repulsive mechanical interactions among the components of the biofilm. We referred as low-depletion effect for elastic-modulus (i) *E_cell–cell_* = *E_eps–eps_* = *E_cell–eps_* = 2 × 10^5^*Pa*,, (ii) moderate depletion effect when *E_cell–cell_* = *E_eps–eps_* = *E_cell–eps_* = 4 × 10^5^*Pa*, and high-depletion effect for (iii) *E_cell–cell_* = *E_eps-eps_* = 4 × 10^5^*Pa*, *E_cell–eps_* = 6 × 10^5^*Pa*. Next, to get a systematic understanding of the role of motility force, we carried out simulations of growing colonies of non-motile (*f_mot_* = 0) and motile (*f_mot_* ≠ 0) cells in presence of differential frictions between the components and for different mechanical interactions. Figure 10 illustrates the different spatial organizations of the growing colonies for different depletion effects in presence of non-adsorbing EPS and varying motility.

**FIG. 10.**
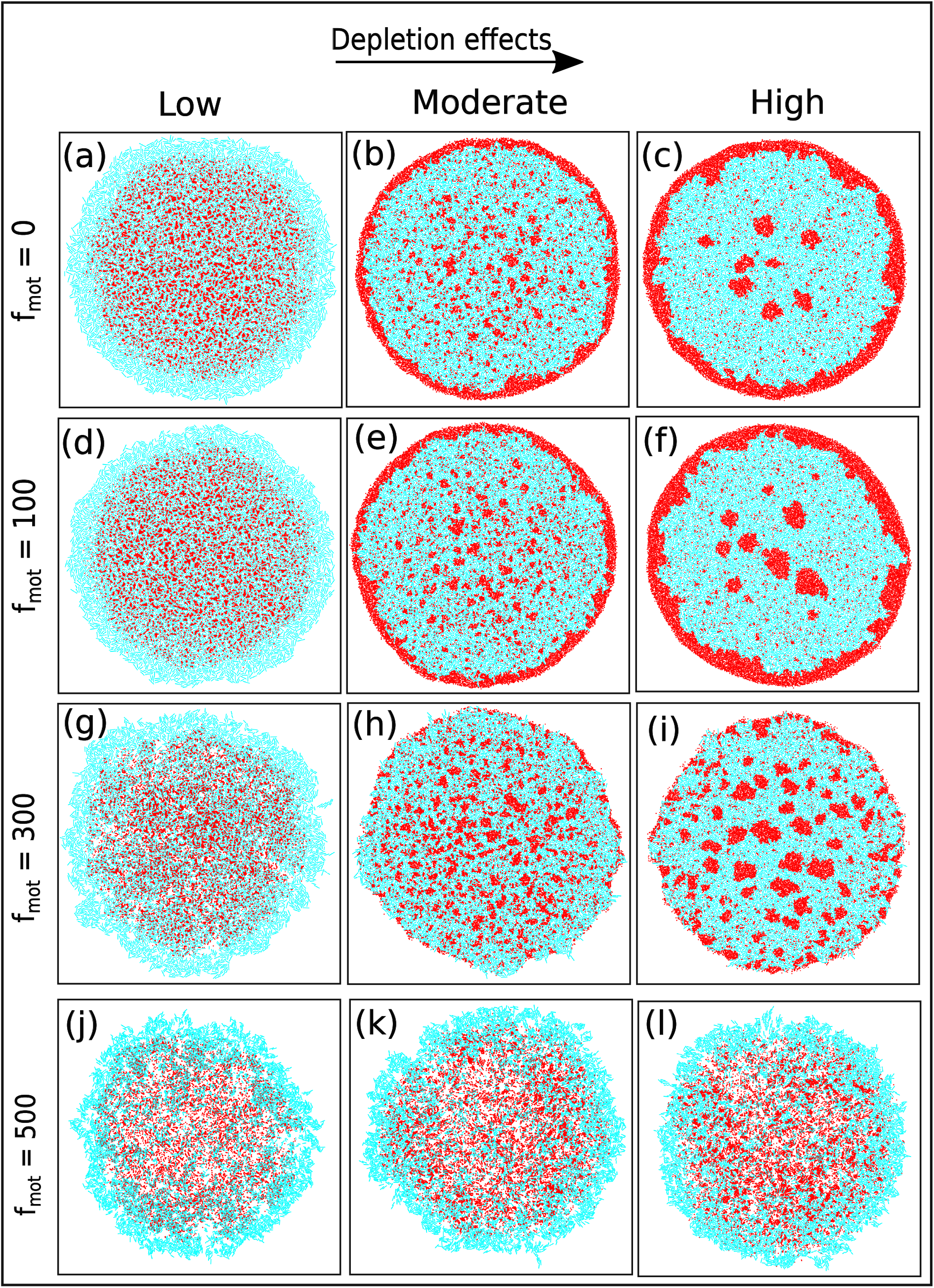
Snapshots of the growing colony for non-motile and motile cells in presence of self-produced non-adsorbing EPS for low, moderate and high depletion effects respectively.

The snapshots of the simulated colonies for low, moderate, and high depletion effects, respectively, are depicted in Figure 10(a), (b), and (c), in the case of non-motile cells (*f_mot_* = 0). A similar spatial organization is also observed in case of growing colonies with motile cells having low motility force such as *f_mot_* = 100*Pa.μm*^2^ as shown in Figure 10(d), (e) and (f). These results reveal that in the presence of differential frictions, motile cells with low self-propulsion forces exhibit a similar spatial morphology as a non-motile variant. Specifically, in the case of a low depletion effect, the interior portion of the colony shows the presence of a mixture of cell and EPS particles surrounded by a rim of only cells at the colony periphery. However, for moderate depletion effects, EPS particles start to move towards the peripheral region of the expanding colony with weak phase segregation in the colony interior rendering a phase-segregated patterned colony as depicted in Figure 10 (b) and (e). On the other hand, for high depletion effects, a large concentration of EPS particles accumulate at the colony periphery developing an annular region of EPS as depicted in Figure 10(c) and (f). Inside the colony, there exists a few phase-segregated regions of EPS particles surrounded by the cells, and these EPS and cells are well phase-separated due to depletion interaction [19, 23, 47, 48, 53]. This unique morphological diversity in the presence of differential frictions, accompanied by mechanically-driven phase segregation, is one of the significant results of the present study. An existing previous experimental study by Srinivasan et al., [25] demonstrated that during the evolution of *Bacillus subtilis* biofilms, matrix production is localized to an annular front, propagating at the periphery. However, the underlying reasons are not clear. Our finding provides one of the plausible mechanisms for which a similar type of spatial morphology might emerge in a growing colony that is just mechanically driven.

To get a deeper insight into the effect of differential mobility of cells and EPS particles in the presence of mechanical interactions, we further performed simulations with increasing values of motility forces such as *f_mot_* = 300*Pa.μm*^2^ and *f_mot_* = 500*Pa.μm*^2^. The corresponding results are depicted in the images in Figure 10 (g) to (l), which demonstrate the snapshots of simulated colonies for low, moderate, and high depletion effects. From these spatial images of growing colonies, it is evident that an increase of *f_mot_* = 300*Pa.μm*^2^ changes the spatial morphology in the presence of differential friction between the cell and EPS. As motility force increases, it reduces the effect caused by differential friction, and for low mechanical interactions, a sparse colony as shown in Figure 10(g) develops. While, in presence of moderate depletion effect, colony starts to phase-segregate (Figure 10(h)) and, for the high-depletion effect, we observe spontaneous phase-segregated spatial organization in a non-equilibrium growing colony as shown in Figure 10(i). Interestingly, we observe a spontaneous phase separation between cells and EPS, giving rise to a patterned colony even in the presence of motility force, albeit moderate. These results infer that there is competition between the particles’ spatial dispersion aided by the self-propulsion of cells and the motion due to their mechanical interactions. To verify this argument, we simulated the morphology of the growing colonies with a high motility force *f_mot_* = 500*Pa.μm*^2^ (Figure 10 (j), (k), and (l)). We find that a large motility force of the cells escalates the cellular motion and forces them to move outside of the colony.

We quantify the spatial morphologies of the colonies belonging to Figure 10 by calculating the radial intensity profiles of cell and EPS particles of the growing microcolonies as demonstrated in Figure 10 for the last few time frames. For each frame, we computed the distance of the particles from the center of the simulation box and made a distribution of these distances. The corresponding count is defined as radial intensity. Figures 11(a) and (b) represent the radial intensity profiles of EPS particles and cells respectively for low depletion interactions, while 11(c) and (d) illustrate the radial intensity of the EPS and cells for the high depletion effects respectively for the low motility forces (*f_mot_* = 100*Pa.μm*^2^). In the case of the low depletion effect, we find that radial intensity profiles are similar for cells and EPS, suggesting a homogeneous mixing of the particles in the colony interior. However, a close look indicates the presence of only a thin rim of cells outside the microcolonies. On the other hand, for the case of high depletion effects, we see a large intensity peak of EPS emerge at a large distance for each time frame which indicates that EPS particles move towards the periphery of the colony in agreement with Figure 10(f). When motility force is reasonably high, we observe that, similar to what happened in the case of the low depletion effect, the radial intensity plots show a mixture of both the particles at the colony interior, followed by a thin rim of cells at the periphery as depicted in figures 11(e) and (f). However, a different micro-colony morphology is observed in case of high-depletion effects (figures 11(g) and (h)). We see several small intensity peaks arising as depicted in Figure 11(g), which represents that some EPS particles are clumped inside the interior of the micro-colony forming small clusters. The effect of differential friction between the two types of particles was introduced by *E/η* ratio term, which behaves like the inverse of diffusion time. For moderate and high depletion effects, this ratio is large compared to the low depletion case, which implies that the diffusion time of EPS is less, which escalates the EPS to move towards peripheral regions. Besides, the dispersion of cells is guided by their self-propulsion ability. Finally, the morphology is controlled by the interplay of motility and differential frictions of the particles.

**FIG. 11.**
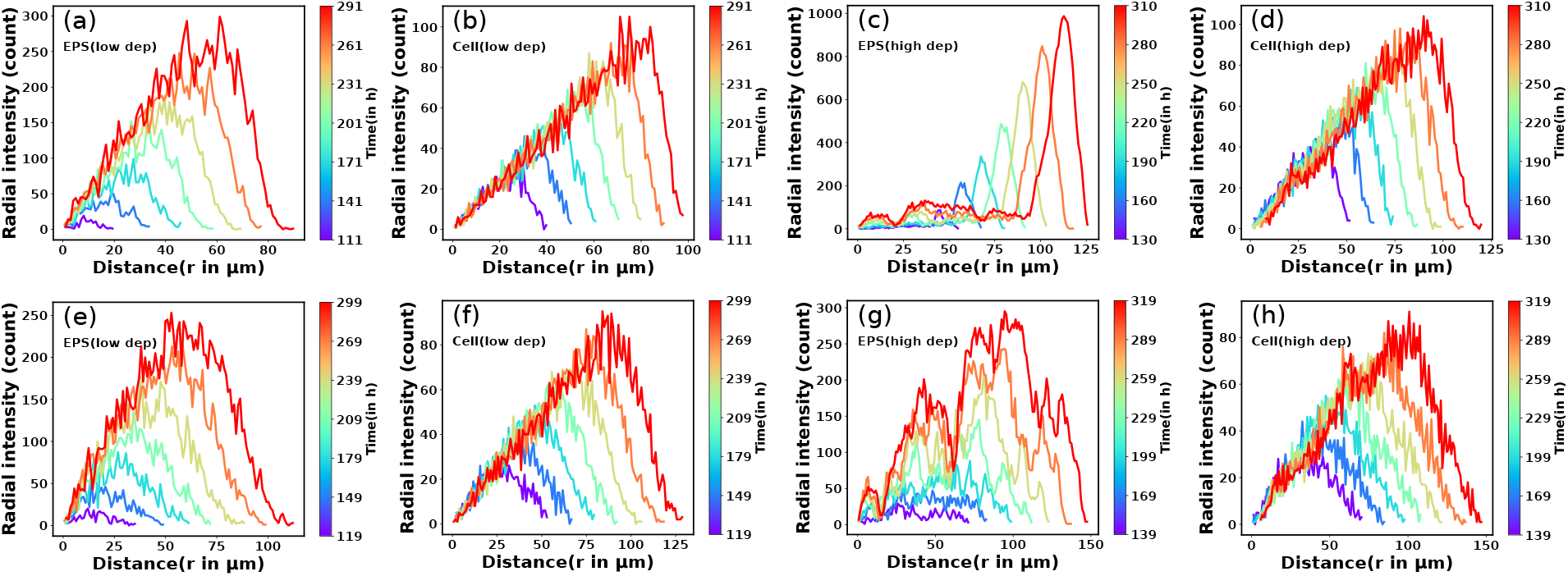
The radial intensity of EPS and cells for different depletion forces as (a) low (EPS), (b) low (cells), (c) high (EPS), and (d) high (cells) respectively with motility force *f_mot_* = 100*Pa.μm*^2^. The radial intensity of EPS and cells for different depletion forces as (e) low (EPS), (f) low (cells), (g) high (EPS), and (h) high (cells) respectively with motility force *f_mot_* = 300*Pa.μm*^2^. Here we have plotted the radial intensity of the last few frames and the color bar denoted the time. For high depletion and low motility force, EPS move to the periphery and form an annular region.

#### Effect of local nutrient accessibility: transition from dendritic to smoother colony periphery

Since cell growth and heterogeneity of EPS production are linked with the local nutrient availability, we will now discuss the effect of initial nutrient concentration, *C*_0_, on the morphological dynamics of non-motile bacteria. We have chosen a slowly diffusive nutrient for the colony growth (*D* = 30*μm*^2^/*h*). In our previous study [30], we have shown that with increasing value of initial nutrient concentration, the colony shows a transition from finger-like to smoother front. The followed-up questions that we ask here are: How does *C*_0_ regulate the colony morphology in the presence of self-produced EPS? To answer this question, we perform a new set of simulations of non-motile cells by varying *C*_0_, keeping all other parameters the same as in Figure 2. Figure 12 demonstrates the snapshots of the growing colony with a variation of *C*_0_(3.0, 7.0,10.0, 20.0) *fg.μm*^3^. We notice that for a small value of *C*_0_ = 3.0f*g.μm*^3^, the colony evolves, forming a dendrite-like structure. With an increase of *C*_0_, colony morphology changes from branched-like to smoother. Cell growth, divisions, and EPS production are low due to the weak nutrient access in a low-nutrient media. As *C*_0_ increases, cell growth, division, and EPS production increase due to increased nutrient access. However, it shows that local nutrient access determines the morphological transition at the colony front, suggesting that EPS interaction does not significantly impact when its production itself is nutrient-dependent.

**FIG. 12.**
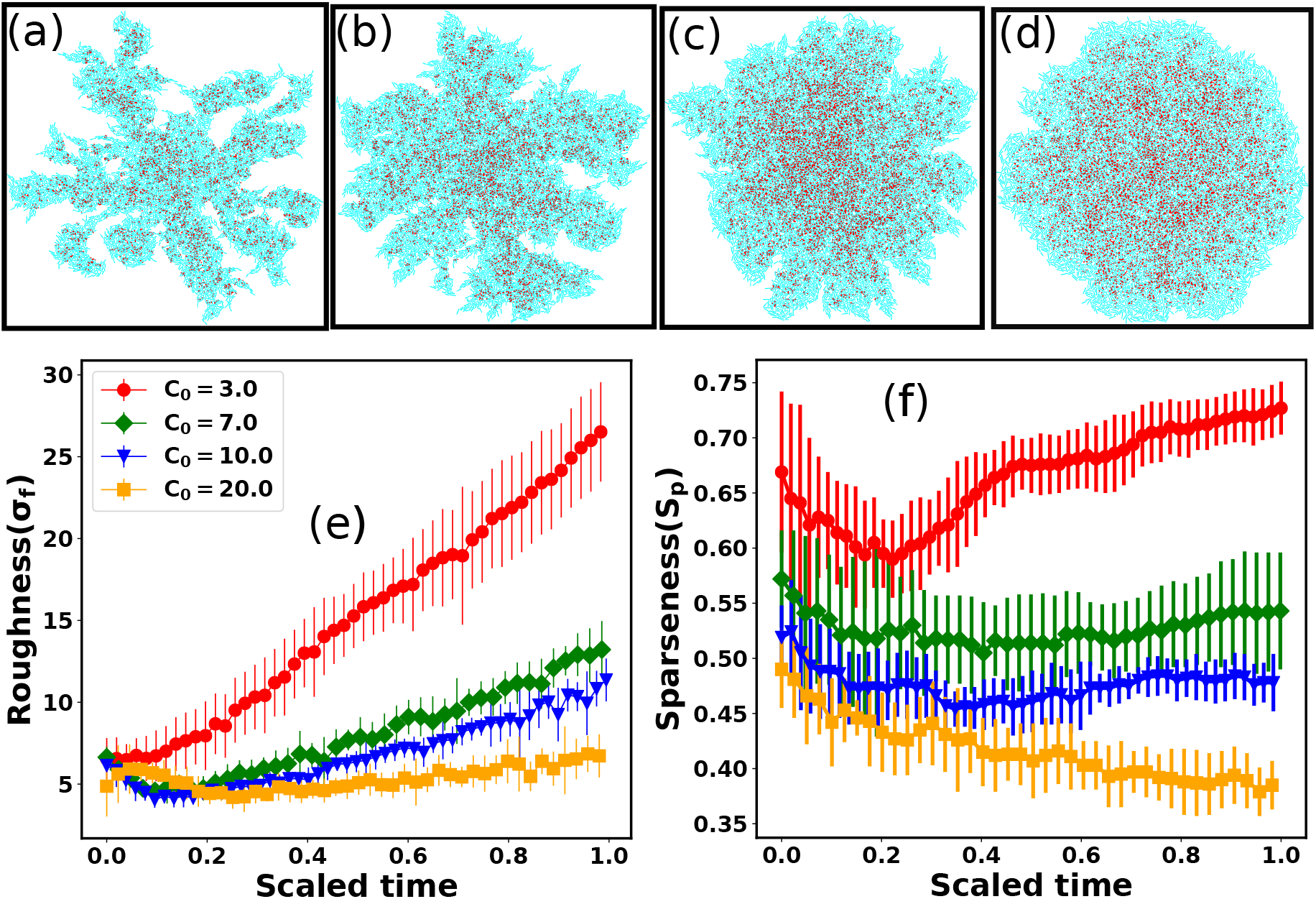
Snapshots of the non-motile growing colony in presence of non-adsorbing EPS for initial nutrient concentration (a) *C*_0_ = 3.0*fg.μm*^3^, (b) *C*_0_ = 7.0*fg.μm*^3^, (c) *C*_0_ = 10.0*fg.μm*^3^, and (d) *C*_0_ = 20.0*fg.μm*^3^ respectively and the nutrient diffusion coefficient *D* = 30*μm*^2^/*h*. Colony becomes dendrite to smoother with increasing values of initial nutrient concentration *C*_0_. Plots of (e) Roughness and (f) sparseness as a function of scaled time for different values of initial nutrient concentration (*C*_0_). Roughness and sparseness, both are higher for smaller values of *C*_0_. For Figure (f) the color coding is the same as the legend of Figure (e). Error bars represent the standard error.

To measure the dendritic to smoother transition at the colony periphery, we estimate the roughness parameter denoted by (*σ_f_*), which is the ensemble average of the standard deviation of the distance of peripheral cells from the center of the simulation box. To calculate *σ_f_*, we divide the colony into specific angular bins (0 – 2*π*) in each time interval, identify cells belonging to those bins, and then estimate the maximum distance from the center of the simulation box. Subsequently, we determine the standard deviation of these maximum distances, referred to as the roughness(*σ_f_*) of the colony periphery. Figure 12(e) shows the time profiles of the roughness parameter for different values of *C*_0_. For low values of *C*_0_, *σ_f_* increases almost linearly. However, for large values of *C*_0_s, *σ_f_* is relatively smaller, which is in agreement with the colony morphology observed as in Figure 12(c), and (d) and saturate at later times.

Finally, to get further insight into the morphological dynamics, we have estimated the sparseness (*S_p_*) of the growing colonies for the different values of *C*_0_. Figure 12(f) illustrates the sparseness as a function of time for increasing values of *C*_0_. We observe that sparseness is inversely proportional to *C*_0_. For colonies growing in lower values of *C*_0_, sparseness is higher. These observations suggest that the morphology at the expanding colony front is majorly regulated by *C*_0_ in addition to the mechanical interactions among the biofilm components.

## IV. SUMMARY AND CONCLUDING REMARKS

Bacterial aggregation, spread, and pattern formation have immense importance for their survival and biological functioning. These features play a crucial role in infections and spreading biofilm formation, and antibiotic resistance. Over the years, experimental and theoretical research have illuminated underlying structural complexities and dynamics in multicellular organizations. Although bacterial cells are rigid and hardly deform under external forces, their multicellular organizations, such as biofilms, are dynamic and active. At the community level, they appeared to be able to generate mechanical forces and respond to the changing environment potentially. Existing studies have established that mechanical forces and physicochemical factors can profoundly influence the spatial morphology and pattern formation in multicellular microbial communities [7–9, 18–21, 29–31, 33]. It is now becoming increasingly relevant to consider mechanical interactions while underpinning the spatiotemporal evolution and dynamics in multicellular systems composed of mesoscopic objects such as bacterial cells.

Here, we have investigated biofilm morphogenesis focusing on the physicochemical properties of self-secreted EPS, cell growth, and motion during microcolony development using a particle/agent-based model from the perspective of soft matter physics. The spatiotemporal dynamics of a growing monolayered microcolony is understood in terms of its primary components, i.e., the rigid rod-like cells and the self-secreted EPS in the media. Furthermore, heterogeneous expression of EPS due to the spatial heterogeneity of the local nutrients set up concentration gradients within the biofilm, which on the other hand, generates mechanical forces relevant for spreading and spatial patterning. Moreover, the properties of EPS profoundly impact the structural integrity and morphological dynamics of the growing micro-colony biofilms.

Our simulation results reveal a dynamic phase transition indicating the presence of coexisting phases of mobile and sessile aggregates of cells and EPS particles during micro-colony morphogenesis. Specifically, in weakly attractive/sticky EPS, we find a dynamics phase transition where cells inside the colony interior form sessile clusters surrounded by motile cells. The cells at the outermost layer remain motile due to the high accessibility of nutrients and less EPS production. We observe the presence of coexisting cells which follow sub-diffusion and super-diffusion simultaneously for a longer lag time scale. Our control simulation of a growing colony in the absence of self-produced EPS does not show such sub-diffusive dynamics even in a longer lag time scale. These observations support that transition from motile to biofilm-like aggregates is mediated by self-secreting sticky EPS. For this type of growing bacterial colonies with motile cells, the self-propulsion force helps the cells to form small aggregates, and sticky EPS increases the adherence of the cells within the clusters. Therefore, the co-existence of two dynamical phases is conciliated by joint ventures of these two properties (cell motility and sticky EPS).

While our model shows that the sticky EPS provides the mechanism of biofilm transition, the spatiotemporal dynamics of growing colonies differ when self-secreted EPS act as depletants in the media because of their nonadsorbing nature on the surface of cells. In this case, we find various spatial morphologies for differential frictions of the particles with the variation of motility forces. For non-motile or weakly motile cells, it shows that a morphological transition occurs for low to high depletion effects exhibiting the presence of EPS at the periphery of the colony. However, for moderate motility forces and differential frictions, the dual effects establish phase-segregated patterned structures across the growing colony, which mimics the results of our earlier work [19] albeit for the non-motile cells. Finally, the colony appears to be sparse and disordered for large motility forces, showing less structural integrity. These observations reveal the difference in the dispersion ability of the biofilm components aided by the motility forces, differential frictions, and depletion effect, which regulates the overall morphology of the growing colonies.

Despite its simplicity, our model provides crucial insights into the spatial morphology and dynamics of a growing biofilm. We find that the combined effects of cell motility, growth-induced stress, and mechanical interactions among the biofilm components regulate the spatial heterogeneity and pattern formation during biofilm morphogenesis. One of the advantages of our model, owing to its simplicity, is that it can predict and provide biophysical intuition for the different behaviors exhibited by a multicellular microbial community for varied conditions.

## ACKNOWLEDGMENTS

All the authors acknowledge Tata Institute of Fundamental Research Hyderabad, and Indian Institute of Science Education and Research Thiruvananthapuram, India for providing the support of computing resources.

