## Supporting Information for "A mechanistic understanding of biofilm morphogenesis: Coexistence of mobile and sessile aggregates and phase-separated patterns"

### 0.1 The Model Details

Each bacteria is spherocylindrical with fixed diameter ($d_0$) and variable-length $L = l + d_0$, where $l$ correspond to the cylindrical length of the particle. The position and orientation of individual cell are decided by spatial coordinate $\vec{r}(x, y)$ and unit vector $\vec{u}(u_x, u_y)$. We inoculate a single bacteria around the center of the simulation box and keep the nutrient concentration fixed to $C_0$ on each grid point. The bacterial colony expands due to the consumption of nutrients which follows a diffusion equation with a sink term as

$$\frac{\partial C}{\partial t} = D\left(\frac{\partial^2 C}{\partial x^2} + \frac{\partial^2 C}{\partial y^2}\right) - k\sum A_i f\left[C\left(x_i, y_i\right)\right] \tag{1}$$

where $x_i$ and $y_i$ are the spatial coordinate, $D$ is the diffusion constant of the nutrients, $A_i = \pi r_0^2 + 2r_0 l_i$ is the area of $i^{th}$ cell, $r_0 = d_0/2$ is the radius of end caps and $l_i$ is the length of the cell. Bacterial cells consumed nutrients at a rate $kf(C)$ per unit biomass density, where $f(C)$ is a monotonically increasing dimensionless function. In our simulation, we consider $f(C) = C/(1 + C)$, a Monod function with a half-saturation constant equal to 1 (in arbitrary units). Each cell grows along their major axis as per the relation $dl_i/dt = \phi \times (A_i/\bar{A}) \times f(c(x_i, y_i))$, where $\phi$ is the linear growth rate of the cell and $\bar{A} = \pi r_0^2 + (3/2) r_0 l_{max}$ is the average cell area [1–3]. Generally once a cell reaches to a critical length $l_{max}$, it splits into two daughter cells, at a rate $k_{div}$, with slightly random orientation compared to the mother cell. This randomness in orientation assimilates the effect of various deformities like the roughness of the agar surface and slight bending of the cells etc. This stochasticity also confirms that the cell will not form a long filament-like structure. But for some species depending on environmental conditions, they show asymmetric division in their growing lifestyle [4]. So in our model, we have also incorporated the asymmetric division in a very simple way. We have defined a quantity, *Division*, by taking the numbers from a Gaussian distribution as *Division* $= exp(-\frac{(l(t)-l_{max})^2}{(w \times l_{max}^2)})$, where $w = 0.0055$, $l(t)$ is the length of the cell, and $l_{max}$ is the maximum length of the cell. In our simulation, we have put a condition that cell will divide if and only if *Division* $>$ rand(), where rand() is the random numbers between 0 to 1. Figure represents Division as a function of cell length. So from the figure it is clear that most of the cells will divide at $l(t) = l_{max}$, but there are also finite probability for dividing the cells at $l(t) > l_{max}$ or $l(t) < l_{max}$.

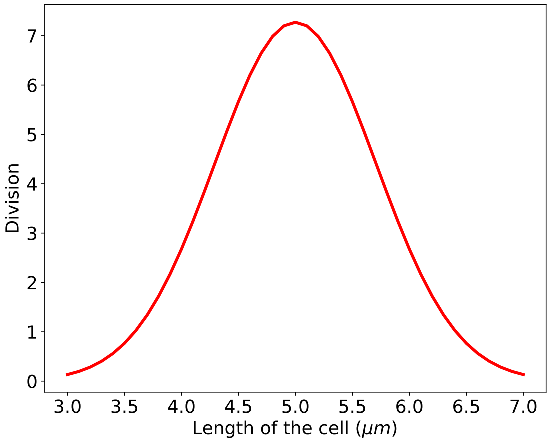


Figure 1: Division as a function of cell length. Most of the cells will divide at $l(t) = l_{max}$, but there are also finite probability for dividing the cells at $l(t) > l_{max}$ or $l(t) < l_{max}$.

It is impossible to determine the conformation of individual EPS on the scale of the bacterial colony. So we have modeled the EPS as spherical particles with a radius equal to the radius of gyration [5, 6]. We have chosen the radius around that half of the radius of the bacterial cells. This radius is larger compared to the actual size of the EPS. So one can think of a single sphere as the assembly of EPS particles. Each cell can produce EPS in the nearby area with certain conditions. These conditions are arbitrated by local nutrient concentration, local cell density, EPS density, etc. The previous experimental studies [7–9] reported that Nutrients depletion in bacterial colonies triggers matrix production. Motivated by these experimental studies, we set a condition that when the local nutrient concentration in each grid point reaches a certain value $C^*$ then the cell can commence producing EPS with a certain rate $k_{eps}$ and place randomly at a small distance from the center of the cell. To avert the excessive production of EPS, we impose a condition to the EPS production by considering that it starts at a location once the local cell area density reaches a certain threshold and stops once the local EPS area density reaches a maximum concentration of EPS [2]. We discretize our simulation box into a certain number of square grids and calculate the cumulative area of cells and EPS that spatially belong to a particular grid point $[x, y]$. Then impose the condition then when the cell's area reaches to a particular threshold (cell[x,y]) then EPS production will start and stop when the EPS area reaches to a particular threshold (EPS[x,y]).

Equation 2 asserts that there are four different interactions acting on the particles (cells and EPS particles). (i) $F_{rf}$ represents the repulsive force between the particles. We assume that mechanical interaction between particles is repulsive, which is in accordance with the Hertzian theory of elastic contact [10,11], if there are spatial overlaps between them. Repulsive force between two spherocylindrical rods is assumed by the force between two spheres, placed along the major axis at such positions that their distance is minimal [2,12]. The repulsive force can be chronicled by the equation: $F_{rf} = Ed_0^{1/2}h^{3/2}$ [1–3,12–15], where $E$ is the elastic modulus of the cells and $h = d_0 - r$ represents the overlap between two interacting cells, where $r$ corresponds to the closest distance of the approach between the two cells. For the repulsive part, EPS particle also follow the same interaction as bacterial cells. (ii) $F_{af}$ represents short range attractive force between cells and EPS particles. Recent experimental study by Vasco M. Worlitzer [16] suggest that only matrix-builders cells can transform from motile to biofilm state. Motivated by this experiment, we have introduced a short range attractive force between cells and EPS particles which mimics that EPS behaves like sticky particles. For attraction interaction, we have used attractive part of Lennard-Jones (L-J) potential i.e $F_{af} = -24\epsilon d_{eff}^6/r^7$ and range of this attraction has taken as $2.5d_{eff}$ where $d_{eff} = (d_0^{cell} + d_0^{eps})/2$. So if there are any overlaps between the particles, they will feel repulsive force and only EPS and cells will sense attractive force if they are in certain cut-off distances. Together with this attractive and repulsive part, we can say that EPS and cells are interacting via full LJ force with soft repulsive part and fixed cutoff distance compare to original LJ potential. (iii) $F_{mf}$ represents the motility force or self propulsion force acting on each cell, along their long axis which mimics the flagella motor speed. (iv) $\zeta$ represents a random force from surrounded medium which is taken from a uniform distribution with a range $-10^{-3}$

to $+10^{-3}$ [15].

$$\dot{\vec{r}} = \frac{1}{\eta L}\vec{F} = \frac{1}{\eta L}(\vec{F}_{rf} + \vec{F}_{af} + \vec{F}_{mf} + \vec{\zeta}) \quad (2)$$

$$\omega = \frac{12}{\eta L^3}\tau \quad (3)$$

### 0.2 MSD calculation

For MSD calculation, we have drawn a circle about the center of the box with a radius $r^* = 60\mu m$ after a certain time $t^* = 268h$. The figure 2 represents the snapshot of the growing colony at that particular time $t^*$ and the circle is denoted in red color. We have tracked only those bacteria which are belonging to this particular radius $r^*$ in that time span $t^*$.

### 0.3 Identification of two type of cells

To classify these two types of cells, we have drawn a circle from the center of mass of the colony with a radius that is slightly smaller compared to the maximum distance of the EPS from the center of mass of the colony. Figure 3 represents a particular snapshot of a growing colony and the cells which are belonging to the circle are named interior cells and cells which are outside of the circle are named exterior cells.

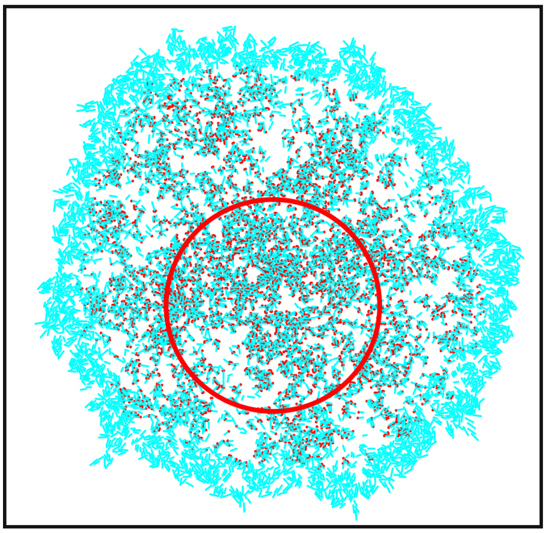

Figure 2: Snapshot of the growing colony at a particular time $t^* = 268h$ with an initial nutrient concentration $C_0 = 3.0fg.\mu m^3$ and self-propulsive force $f_{mot} = 500Pa.\mu m^2$. We have tracked only those bacteria which are belonging to this particular radius $r^*$ (red circle) in that time span $t^*$.

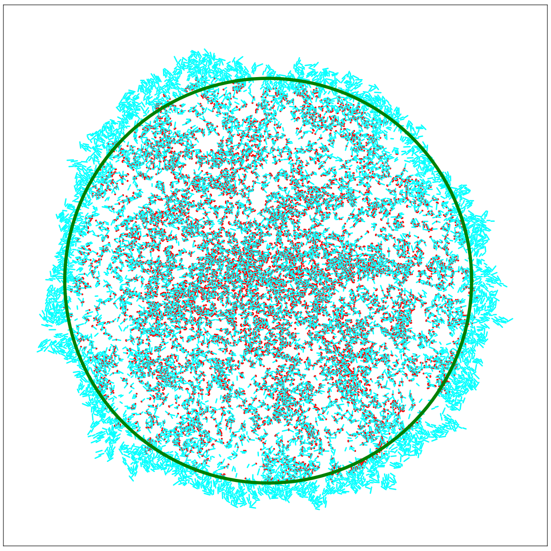

Figure 3: Snapshot of the growing colony at a particular time. The cells which are belonging to the circle are named interior cells and cells which are outside of the circle are named exterior cells.

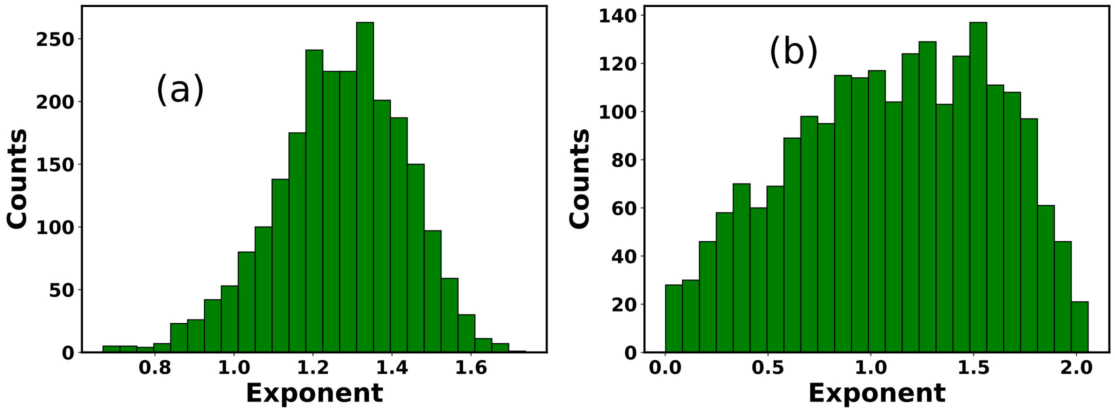


Figure S1: Distribution of MSD exponents of cells in presence of sticky EPS for initial nutrient concentration $C_0 = 10.0 fg.\mu m^3$ with two different lag time (a) small ($\tau_1$) and (b) large ($\tau_2$) respectively. For both cases more cells are showing sub-diffusion in two time scale compare to initial nutrient concentration $C_0 = 3.0 fg.\mu m^3$.

Table S1: **Parameters and constants used in our agent-based model**

| Parameter | Symbol | Simulations |
|---|---|---|
| Maximum length | $l_{\max}$ | 5.0 $\mu$m |
| Diameter of cell | $d_o$ | 1.0 $\mu$m |
| Diameter of EPS particle | $d_{\text{eps}}$ | 0.5 $\mu$m |
| Linear growth rate | $\phi$ | 3.5 $\mu$m/h |
| Cell division rate | $k_{\text{div}}$ | 0.1 /h |
| EPS production rate | $k_{\text{eps}}$ | 1.0 /h |
| Elastic modulus (cell and EPS) | E | $2 \times 10^5$ Pa |
| Friction coefficient (cell) | $\eta$ | 200 $\text{Pa} \cdot \text{h}$ |
| Friction coefficient (EPS) | $\eta_{\text{eps}}$ | 200 $\text{Pa} \cdot \text{h}$ |
| Nutrient concentration | $C_0$ | 3.0, 10.0, 20.0, and 30.0 $\text{fg} \cdot \mu\text{m}^3$ |
| Nutrient consumption rate | k | 4.0 /h |
| Diffusion rate of nutrient | D | 300 $\mu\text{m}^2$/h |
| Threshold area-density of cell | Cell [x, y] | 8.0 $\mu\text{m}^2$ |
| Threshold area-density of EPS | EPS[x, y] | 0.3 $\mu\text{m}^2$ |
| Concentration cut off for EPS production | $C^*$ | 0.006 $\text{fg} \cdot \mu\text{m}^3$ |
| Motility force | $f_{mot}$ | 100, 300, 500, and 700 $Pa.\mu m^2$ |
| Strength of attraction | $\epsilon$ | 18.0 $Pa.\mu m^3$ |
